# Effects of Post-Activation Potentiation induced by a plyometric protocol on deceleration performance

**DOI:** 10.1101/766097

**Authors:** Gianmarco Ciocca, Harald Tschan, Antonio Tessitore

**Affiliations:** University of Vienna, Centre for Sports Science and University Sports, Vienna, Austria; University of Rome “Foro Italico”, Department of Movement, Human and Health Sciences, Rome, Italy

**Keywords:** braking, potentiating, change of direction, warm-up, jumping, bounding, unilateral, power

## Abstract

Post-Activation Potentiation is a phenomenon by which muscular performance characteristics are acutely enhanced as a result of their previous contractile actions. It has been shown how Post-Activation Potentiation, which is usually evoked through heavy resistance exercise, has the potential to improve many different power performances, such as sprinting and jumping. Due to an easier applicability, some studies explored the potential of plyometric muscular actions to evoke the effects of Post-Activation Potentiation. Despite some findings on acceleration running performance, to the authors’ best knowledge, no studies investigated the effects of Post-Activation Potentiation on deceleration performance, which is a key factor in sports involving change of directions. Therefore, the aim of this study is to investigate the influence of a plyometric exercise protocol to a subsequent deceleration running performance. University soccer players (n = 18) performed 7 deceleration trials: at baseline and after ∼ 15 seconds, 2, 4, 8, 12 and 16 minutes a walking control condition (C) or 3 sets of 10 repetitions of alternate-leg bounding (plyometric, P). Results show that no significant differences were found at any of the trials of the control condition (C) in comparison to the relative baseline. In the plyometric condition (P), the deceleration performance executed 2 minutes after the plyometric activity resulted significantly faster compared to the relative baseline (p = 0.042; ES = 0.86, large effect; % of improvement = 4.13 %). Therefore, the main findings of this study showed that a plyometric exercise has the potential to improve a subsequent running deceleration performance in soccer players, if an adequate recovery between these activities is provided to the players. These findings encourage further future investigations about the possible potentiating effects of plyometric activities on more complex actions like changes of direction and agility.

## Introduction

In many sports, the act of rapidly slowing the body (deceleration) is a key factor to the success of the movement [1]. The deceleration plays an important role for the players’ movement patterns in many team sports, such as soccer [2], field hockey [3], rugby [4] and others. In these team sports, running requires changes in velocity through acceleration and deceleration [5]. In fact, during a game, the ability to rapidly change velocity and direction is a key factor for the outcome of many technical actions [6], such as regaining the ball after a loss of possession and evading opposition tackles [7]. Indeed, according to the agility’s deterministic model proposed by Sheppard and Young [8], the player is required to suddenly adapt his own movement to that of his opponent and the current situation.

During a soccer match, the number of accelerations and decelerations and the mean and maximum distance covered during these actions vary according to the intensity of the movements. Mara et al., [7] in a study on elite female soccer showed as players performed a mean of 430 decelerations per match, with differences in intensity according to their position and time period of the match. In this study, both mean and maximum time of interval between two consecutive decelerations was lowest during the first 15 minutes of play compared to all the other 15-minute periods, and this may be attributed to a combination of factors including physical and mental fatigue, game specific factors or strategies [7]. Furthermore, Dalen et al. [9] also demonstrated that decelerations contributed to 5 – 7 % of the total player load during a match, and not considering their energy cost could lead to an underestimation of the player’s match total load [10]. For this reason, it has been suggested how the use of only speed and distance variables to assess the physical demands of soccer players may be limited. In fact, high intensity activities such as jumping, accelerate or decelerate may be classified in the low-speed locomotor category, although they represent a high physical strain for the player [9].

The primary muscles used to decelerate in running actions are the quadriceps and gastrocnemius, working through eccentric muscle actions to absorb and disperse the impact forces, which can be very high if the time available to absorb them is small [1]. Consequently, it has been suggested that decelerating might require a higher decrease in the acceleration forces rather than an increase in the deceleration ones [5].

Post-Activation Potentiation (PAP) is a frequently studied subject in sport scientific research and it has been defined as a phenomenon in which the contractile history of skeletal muscle may facilitate the volitional production of force [11]. PAP refers to the phenomena by which muscular performance characteristics are acutely enhanced as a result of their contractile history [12]. Several authors have demonstrated how PAP can acutely increase muscular power and, consequently, performance [13]. PAP is induced by a voluntary conditioning contraction (CC), performed typically at maximal or near-maximal intensity, and has consistently been shown to increase peak force and, especially, rate of force development (RFD) during subsequent twitch contractions, enhancing the mechanical power (Force×Velocity) and then the sport performances largely determined by it [11, 12]. The hypothesized mechanisms responsible for PAP are the phosphorylation of myosin regulatory light chain (RLC), an increase in the recruitment of higher order motor units and a decrease in muscular pennation angle [12]. After a conditioning activity, mechanisms of muscular fatigue and potentiation (PAP) coexist, and the subsequent power output and performance depend on the balance between these 2 factors [13]. Although twitch studies have reported maximal PAP immediately after a CC, fatigue is also present early on diminishing or unchanging the performance of subsequent voluntary activity. [12]. However, fatigue subsides at a faster rate than PAP, and potentiation of performance can be realized at some point during the recovery period [12]. In exploiting PAP to enhance performance, according to Sale [14], two dilemmas must be taken into account: a more intense and prolonged conditioning activity may activate the PAP mechanisms to a greater extent, but it also produces greater fatigue. Then, a longer recovery period between the end of the conditioning activity and the beginning of the performance may lead to a greater recovery from fatigue, but also to a greater decay of the PAP [14].

There is a combination of several variables influencing the magnitude of PAP and its relationship with fatigue: volume, intensity and type of the CC performed, subject characteristics such as training status and fibre-type distribution, type of the activity performed after the CC, rest period length and others [12, 13, 15]. By the meta-analysis of Wilson et al. [13], it has been shown that moderate rest period lengths (7-10 minutes) may elicit the best power output after a conditioning activity, but more trained people may benefit of a shorter recovery time (3-7 minutes) to have the greatest PAP effects. It has also been shown that both isometric and dynamic muscle actions may elicit PAP, though through different mechanisms [12, 13]. In addition to acute enhancing-performance application (for example, in the warm up prior to a competition), PAP has been investigated for his possible application in a long-term training scenario, the such-called complex training. Complex training has been defined as a training strategy that involves the execution of a heavy resistance exercise prior to perform a plyometric explosive movement with similar biomechanical characteristics, in order to gain superior chronic neuromuscular power adaptations [11, 12, 15, 16]. However, the effectiveness of complex training compared to other training modalities is yet to be properly determined [11, 16, 17].

In addition, several studies investigated the effects of PAP on subsequent power performances. Saez Saez de Villarreal et al. [18] showed that high-intensity dynamic loading (80-95 % 1 Repetition Maximum) and a specific volleyball warm-up protocol including various plyometric exercises both enhanced the following jumping performance in volleyball players. Boullosa et al. [19] reported that introducing recovery intervals between half squat repetitions (cluster set) seems to allow a more rapid improvement of jump height compared with a set without recovery period, probably due to a better fatigue-potentiation relationship. Bevan et al. [20] reported that 3 back squat repetitions at 91 % 1 Repetition Maximum (1 RM) improved sprint ability (over 5m and 10m) of professional rugby players, providing adequate and individualized recovery between the conditioning activity and the subsequent sprint activity (the majority of subjects performed their best sprint times at 8 minutes after the preload stimulus). Many authors have also suggested that ballistic (plyometric) activities may be used to elicit PAP [17, 21], also because of their kinematic similarities to subsequent explosive sport activities [12]. The plyometric training (PLY), using the stretch-shortening cycle muscle action, is frequently used as a training method to improve neuromuscular function and to improve both explosive and endurance performances, and it is considered as a bridge between strength and speed, therefore power [22]. Maloney et al. [17] reported that, using plyometric conditioning activities, the recovery time needed to observe the greatest PAP effects may be lower than would be with the use of heavy resistance exercise, suggesting that plyometric activities may elicit less fatigue. In fact, with the use of plyometric conditioning activities, recovery durations of 1-6 minutes have been shown to successfully elicit PAP in many cases [17]. Moreover, in the same review Maloney et al. [17] also reported that performance improvements induced by plyometric exercises-based PAP range from 2 to 5 %, like those induced by heavy resistance exercise. The effects of PAP elicited by plyometric exercises (3 sets of 10 alternate-leg bounds) on the subsequent sprint acceleration performance (over 20 m, with a split at 10 m) have been investigated by Turner et al. [21]. In this study, the sprint acceleration performance was evaluated at baseline, 15 seconds, 2, 4, 8, 12 and 16 minutes after the intervention protocols (control, plyometric or weighted plyometric). Results showed that 10-m sprint performance of plyometric condition was enhanced at 4 minutes recovery relative to its own baseline and to the same-time performance of the control condition. The 20-m performance of plyometric condition was improved following four minutes of recovery. Results also suggested that, with the addition of a weighted vest, it may be firstly elicited a greater fatigue, but a greater PAP-induced enhanced performance later.

However, considering that there are many biomechanical differences between acceleration and deceleration in sport [1], in view of the above-mentioned aspects and given that decelerations are just as common as accelerations in soccer [9, 23] it could be worthy to study the effect of a PAP protocol on the motor ability to decelerate. Therefore, the aim of this study was to investigate whether the submission of a plyometric protocol presented by Turner et al. [21] may elicit the PAP and thus improve the subsequent deceleration performance in soccer players.

## Materials and Methods

### Experimental approach to the problem

To investigate the effects of a plyometric protocol on subsequent deceleration performance, the protocol provided by Turner et al. [21] has been used to assess its effects on acceleration performance. The only difference with the experimental design of Turner et al. [21] has been that in our current study the plyometric weighted condition was not applied. However, accordingly with the abovementioned authors [21] a full control condition that did not include the execution of the preload stimulus was added to the research design to avoid the possible covariate additive effects (fatiguing or potentiating) from repeated maximal performance efforts (deceleration tests).

The subjects had to complete a standardized warm-up, followed by a baseline 10-m deceleration assessment, and then they had to execute either the walking control condition (C) or the preload stimulus of the plyometric condition (P). After completing one of the two conditions, the participants’ deceleration performances were re-evaluated at 15 seconds, 2, 4, 8, 12 and 16 minutes following the respective condition (C or P).

### Subjects

Eighteen (18) male university student soccer players (age 22 ± 2 years) voluntarily took part in this study after giving their informed consent. All the subjects were members of the University of Rome “Foro Italico” soccer team and they were engaged in 3 trainings and 1 official match weekly. The subjects had no previous muscular injures in the last 60 days before the test days. All the subjects had a multi-year experience in soccer and plyometric training. An informed consent form based on Declaration of Helsinki ethical principles for medical research involving human subjects was read and signed prior to testing and informed consent was obtained from all participants before enrollment and testing. The study was approved by the internal review board of the University of Rome Foro Italico, since the sample involved only players (all students) of the university soccer team.

### Procedures

Before the two experimental sessions (day 1 and 2) where participants were submitted the P and C conditions, respectively, they attended a familiarization session in which they practiced both protocols for the deceleration test and plyometric exercise. All sessions were planned on the same artificial-grass soccer field where participants had trained during the season. Participants were instructed to minimize the foot contact time and maximize the horizontal (rather than vertical) impulse during the execution of plyometric bounds, according to Turner et al. [21]. During the two experimental trials, participants performed the same standardized warm-up used in the study by Turner et al. [21], which consisted of jogging (∼ 3 minutes), dynamic stretching exercises for the musculature primarily involved in the subsequent explosive activities (∼ 10 minutes), increasing-intensity sprints and decelerations (decelerations were the only addition in the warm-up protocol) for ∼ 5 minutes. After an active recovery of 2 minutes, participants performed the baseline deceleration test, followed by a further active recovery period of 2 minutes that preceded the execution of both P (in the first session day) and C (in the second session day) conditions. Finally, participants performed the deceleration test at 15 seconds, 2, 4, 8, 12 and 16 minutes after the respective condition (P and C). (Fig 1).

**Fig 1.**
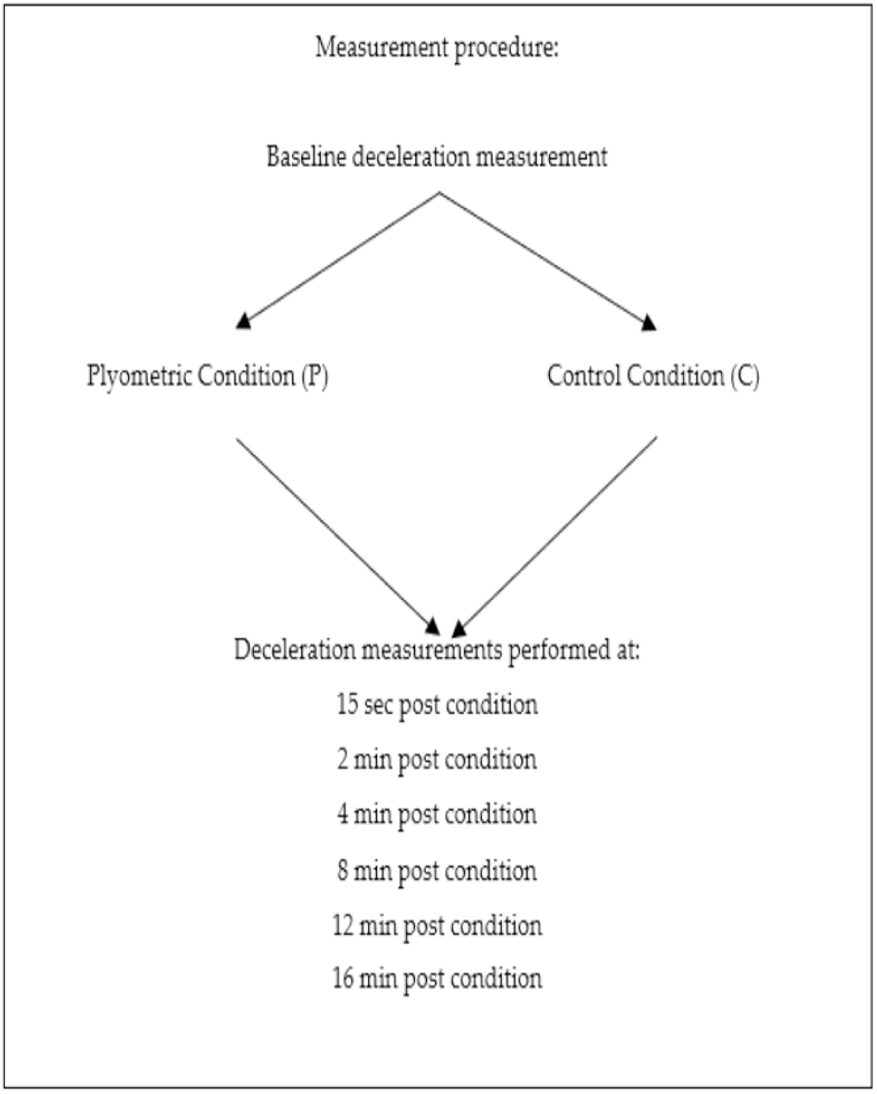
Schematic representation of the measurement procedure.

### Interventions

In the plyometric condition (P), participants performed 3 sets of 10 alternate-leg bounds (5 contacts per leg per set). Once completed the first set, they were instructed to walk back to the starting position and performing the further two sets in the similar manner. Each set (bounds plus recovery to the starting position) lasted for ∼ 25 seconds, as prescribed previously in the protocol by Turner et al.’s [21]. In the control condition (C), participants were instructed to continuously walk for ∼ 75 seconds in order to minimize losses in body temperature relative to the P condition, which lasted about the same amount of time (∼75 seconds).

### Measurements

The ability to quickly decelerate was evaluated by measuring the time to perform a task of acceleration and deceleration over a distance of 10 m followed by a deceleration zone of 30 cm (Fig 2), by means of infrared timing gates (Polifermo, Microgate, Italy) positioned at 0 m (start) and 10 m (finish). Participants had to perform a maximal acceleration, then decelerate and quick stop into a 30-cm deceleration zone placed beyond the finish line at 10 m, as in the protocol by Tessitore et al. [24]. A member of the research staff provided verbal encouragement to the participants in order to assure their maximal effort until the finish line.

**Fig 2.**
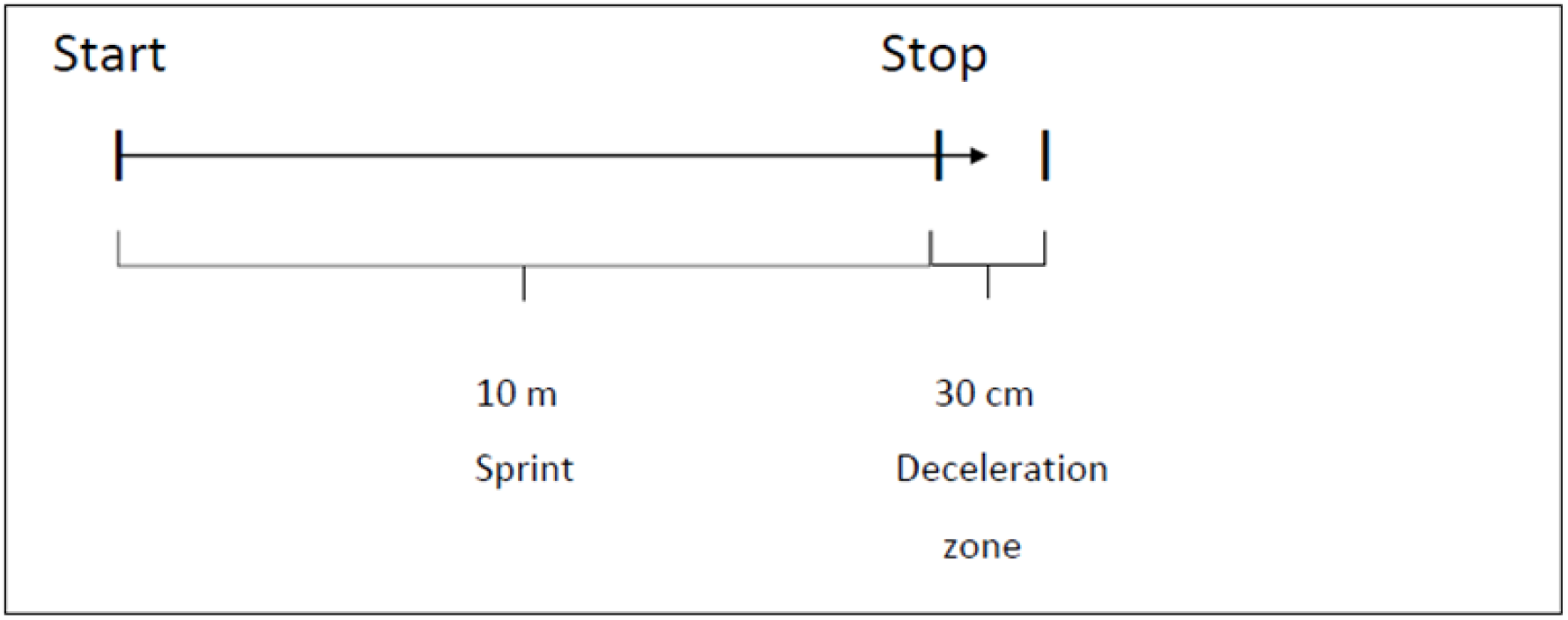
Deceleration test, as proposed by Tessitore et al. [24].

### Statistical analysis

Statistical analyses were performed using SPSS software (Version 23.0, SPSS Inc., Chicago, IL, USA). Data are presented as mean ± SD. The level of significance was set at *p ≤ 0.05.* Before the statistical analysis, the Shapiro-Wilk test of normality was used, showing that data were normally distributed. Two-way (2 × 7) repeated measures analyses of variance (within-subject factors: condition [C, P] × time [baseline, 15s, 2, 4, 8, 12 and 16 minutes]) were used. Mauchly’s test was consulted and Greenhouse-Geisser correction was applied if sphericity was violated. Pairwise comparisons (t-tests) were undertaken for significant main effects for the 6 comparisons of time made for each condition (P and C) to determine differences between the baseline deceleration performance and each of the subsequent intervals’ performances. Cohen effect size (ES) calculations were used with thresholds set at ≤ 0.2, 0.21–0.5, 0.51–0.8, and ≥ 0.8 for trivial, small, moderate, and large magnitudes of effect, respectively.

## Results

The times (mean ± SD) occurred to perform the 10-m deceleration test at different time periods (from baseline to the 16^th^ minute) for both P and C experimental conditions are shown in Figs 3 and 4, respectively. The ANOVA for repeated measures showed a significant difference (p < 0.001) between conditions (P and C) and recovery duration, respectively (15 s, 2, 4, 8, 12 and 16 minutes). A significant difference (p < 0.05) was also found for the interaction condition × recovery duration. The post-hoc analysis for P demonstrated that the deceleration test performed at 2 minutes after the preload stimulus was significantly faster than the baseline one (p = 0.042; ES = 0.86, large effect; Δ time = - 4.13 %), while no significant differences were found for C condition. A comparison between the two experimental conditions (P and C) is also shown in Fig 5. The plain times of performance (mean ± SD) and differences expressed in percentage (%) between baseline and the other time-period (15s, 2, 4, 8, 12 and 16 minutes after the preload stimulus) for P condition are shown in Table 1.

**Table 1.** Mean ± SD and differences in time expressed in percent (Δ %) between deceleration test times and the relative baseline in P condition. * Significant difference from baseline (p ≤ 0.05). Note: lower values mean improvement of performance compared to baseline measurement.

| Deceleration tests at different timing periods | Time for deceleration test (sec) | Percent differences compared to baseline measurement ($\Delta$ %) |
| --- | --- | --- |
| Baseline measurement | 2.34 ± 0.11 | - |
| 15 sec post-condition measurement | 2.35 ± 0.13 | + 0.31 |
| 2 min post-condition measurement * | 2.24 ± 0.12 | - 4.13 |
| 4 min post-condition measurement | 2.27 ± 0.12 | -3.16 |
| 8 min post-condition measurement | 2.30 ± 0.14 | - 1.85 |
| 12 min post-condition measurement | 2.31 ± 0.11 | - 1.45 |
| 16 min post-condition measurement | 2.33 ± 0.10 | - 0.5 |

**Fig 3.**
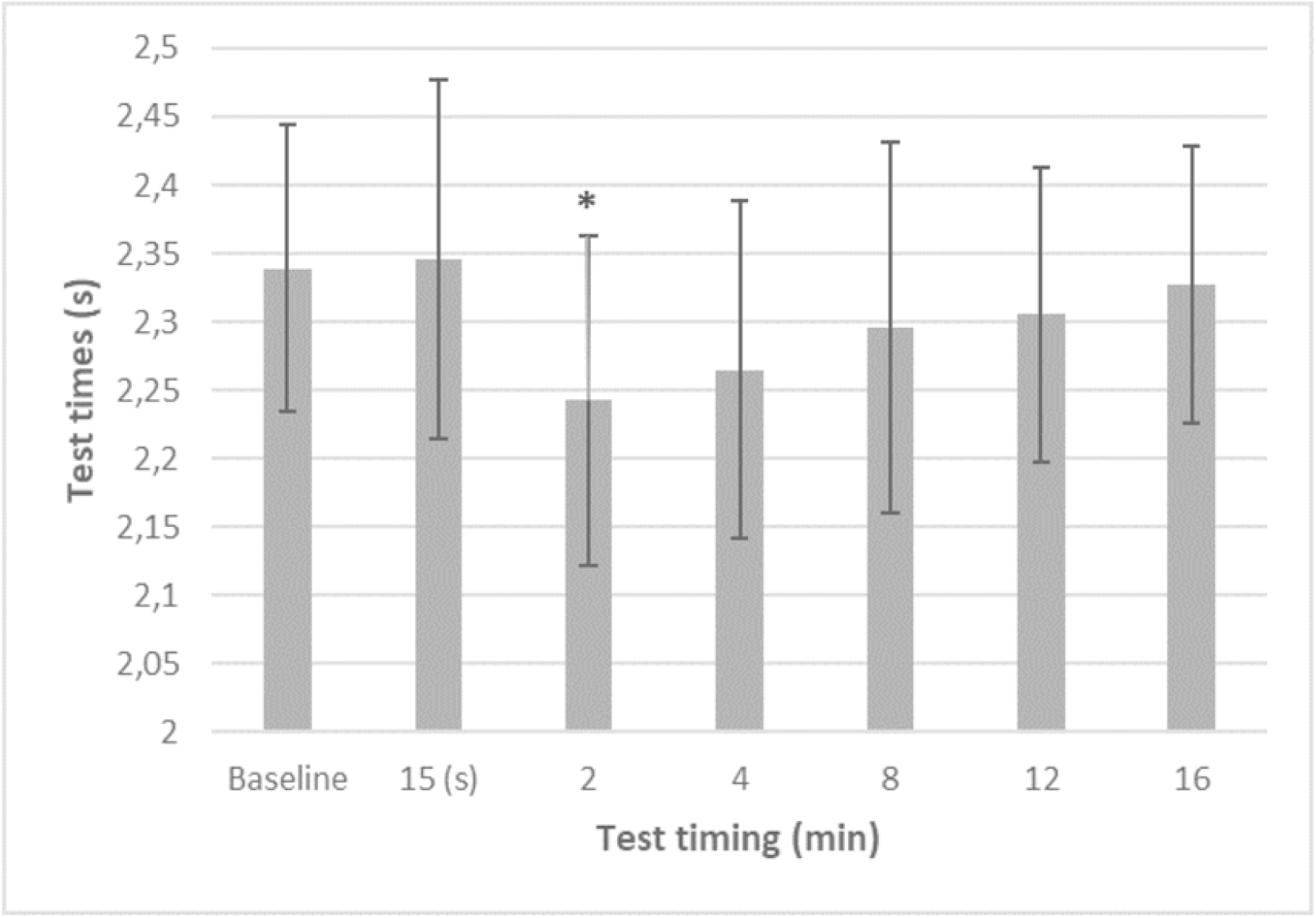
Mean ± SD of deceleration test times of the Plyometric condition (P). * Significant difference from baseline (p ≤ 0.05). Note: lower values mean improvement of performance compared to baseline measurement.

**Fig 4.**
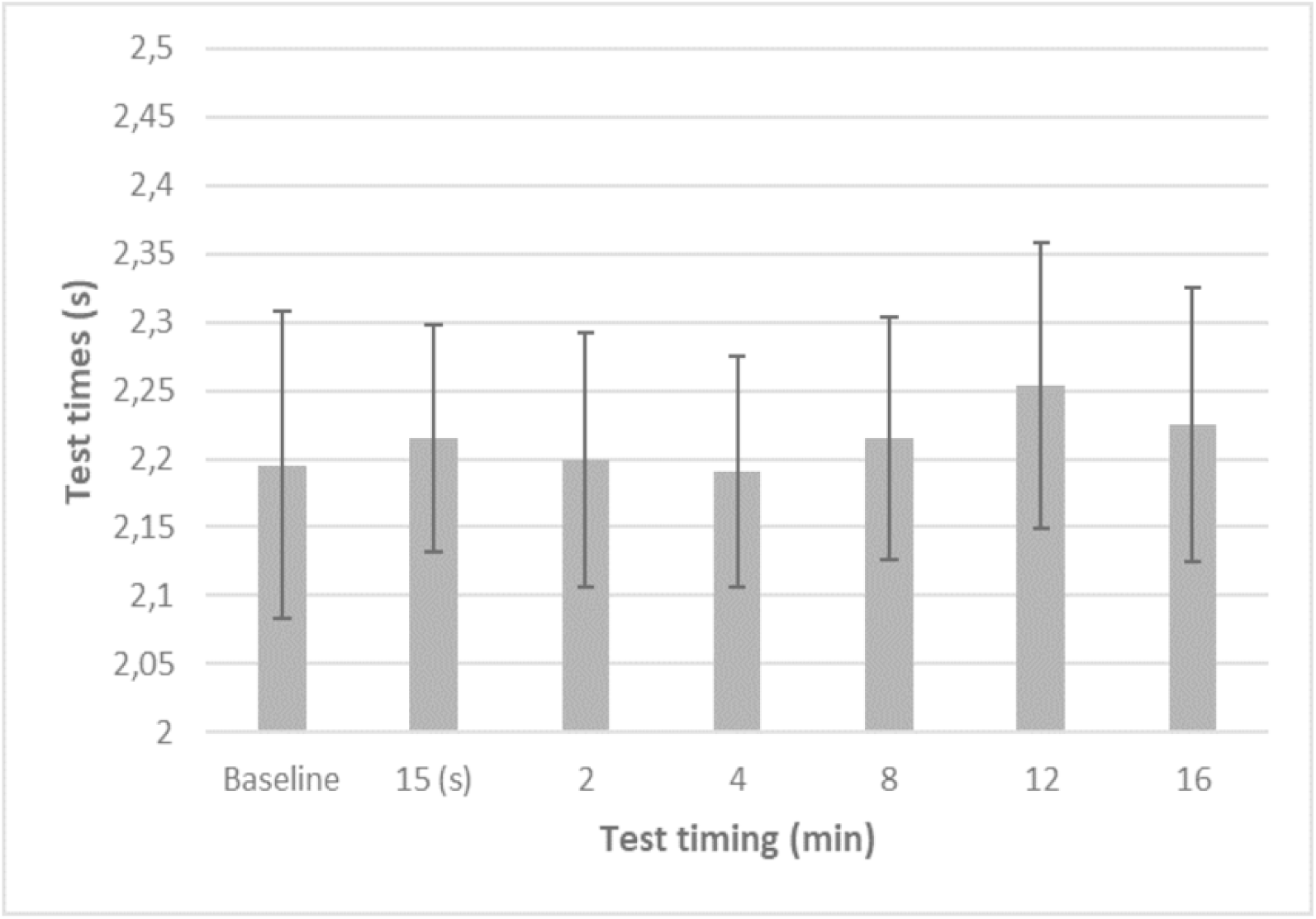
Mean ± SD of deceleration test times of the Control condition (C).

**Fig 5.**
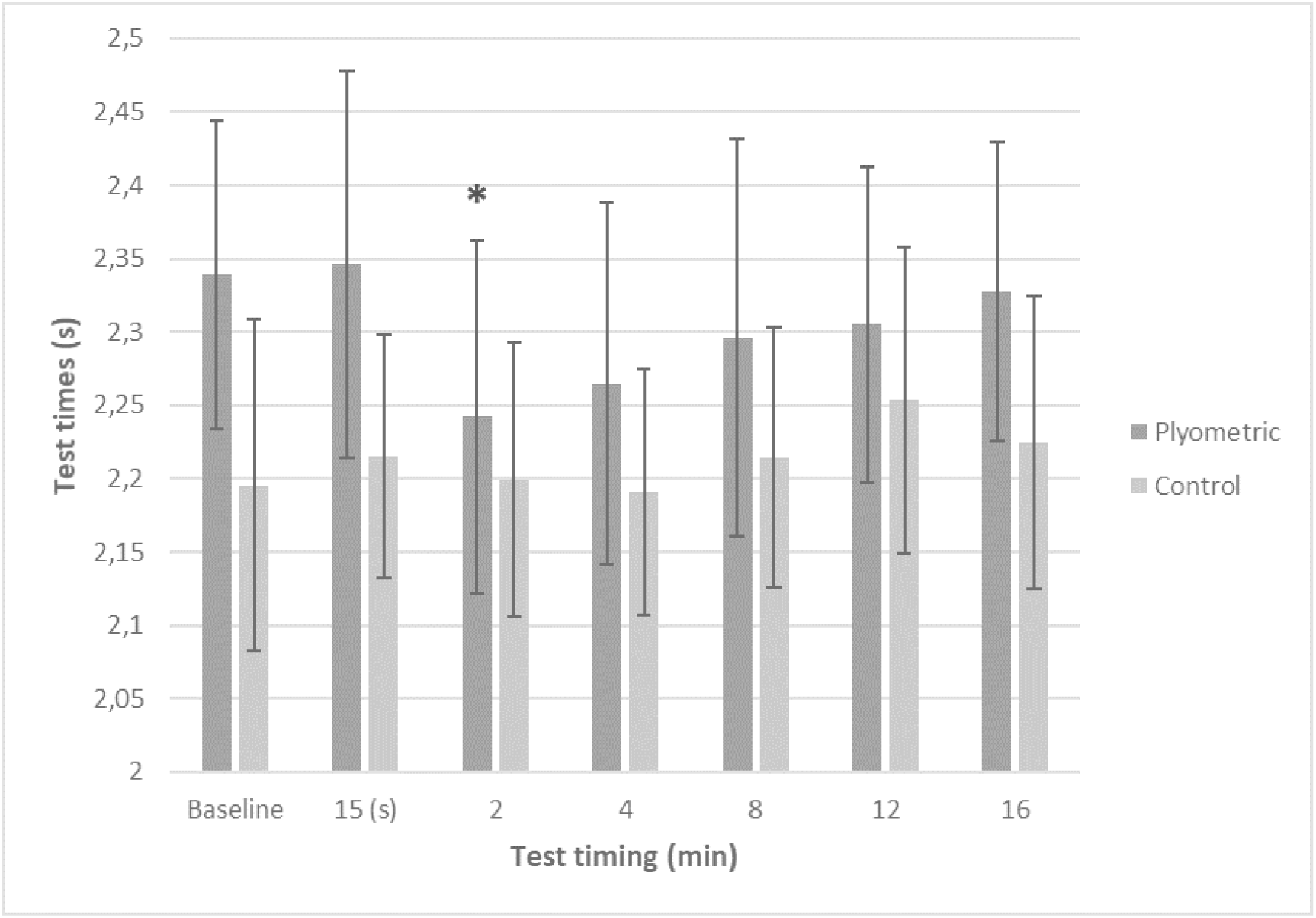
Mean ± SD of deceleration test times of the Plyometric (P) and Control (C) conditions. * Significant difference from the relative baseline (p ≤ 0.05).

## Discussion

To the authors’ best knowledge, this is the first study to investigate the effectiveness of a PAP evoked by a plyometric protocol [21] on the ability to decelerate in soccer players.

The main finding of this study is that the PAP evoked a significant improvement of the players’ performance to decelerate at 2 minutes from its execution, which is also supported by the fact that no significant differences were found between the baseline and the following deceleration performances when the C condition was applied. In terms of percentage, the trial of deceleration performed after 2 minutes from the plyometric preload stimulus improved by 4% compared to its relative baseline, in line with the findings of Maloney et al. [17] who stated that the PAP improvements typically range from 2 to 5 %. Nevertheless, even if the statistical significance has been reached only 2 minutes after the plyometric stimulus (p = 0.042; ES = 0.86, large effect), performance improvements obtained after 4 minutes (- 3.16 %) and 8 minutes (- 1.85 %) should not be underestimated. In fact, Stone et al. [25] reported that differences between first and fourth place in most sports during recent Olympic Games be typically less than 1.5 % [25, from unpublished data].

At present, to our knowledge there are no studies that have examined the effects of PAP on deceleration performance. Turner et al. [21], using the same plyometric protocol as in current study to assess accelerations found improvements of performance at 4 minutes from the preload stimulus and not after only 2 minutes, as in our case. According to Wilson et al. [13], this fact might suggest that our participants could have experienced less fatigue performing the plyometric protocol compared to Turner’s participants. The reason of such fatigued condition could be due to their probable higher training status, fatigue resistance and a certain ability to decelerate. The latter, can also be ascribed to the fact that a part of soccer players’ training load is represented by decelerations themselves, so they could be likely more familiar to perform them.

The results of this study also demonstrate that a plyometric protocol can improve deceleration performance eliciting PAP effects if an adequate recovery time is provided to athletes. Indeed, the deceleration test executed after 15 seconds from the plyometric stimulus was not different compared to its relative baseline, with only a slight decrease in performance (Δ time = + 0.31 %). As evidenced by literature, fatigue and PAP coexist, and subsequent performance depend on the balance between these 2 factors [13]. Our findings suggest that the optimal window with the most effective PAP/fatigue ratio and in which the deceleration performance should be executed is at 2 minutes from the preload stimulus. While the meta-analysis by Wilson et al. [13], recommends recovery periods of 7-10 minutes for optimizing performance enhancement following pre-conditioning our results are more in line with those from Maloney et al. [17] who recommends a rest of 1-6 minutes after plyometric activities as preload stimulus. A possible explanation could be, that the plyometric activity causes less fatigue compared to heavy loading close to 1 RM (would be great if you could find scientific proof or a physiological explanation for that based on neuro-mechanic or energetic research).

According to Tillin and Bishop [12], who proposed that the less the fatigue produced by the preload stimulus, the earlier the PAP enhancing-effects would be seen, our findings suggest that plyometric activities (i.e. alternate-leg bounds) may induce less fatigue than heavy resistance exercises. Supporting this assumption, Turner et al. [21] found improvements in the acceleration performance of university students at 4 minutes from the plyometric conditioning activity, while Bevan et al. showed the best improvements in acceleration performance at 8 minutes following a heavy resistance protocol, although in élite rugby players. This discrepancy in the optimal recovery time between Turner and Bevan’s studies could have even been greater if the participants’ characteristics were more similar, since it has been seen that more trained athletes (such as Bevan study’s rugby players) may benefit from a PAP earlier than less trained athletes, based on their better fatigue resistance [13].

Moreover, the differences found between the current study and the one of Turner et al. [21] concerning the optimal recovery period (2 versus 4 minutes, respectively) may rely also on the fact that decelerations and accelerations have several biomechanical differences [1]. In fact, it has been seen that the type of activity to perform following the pre-conditioning exercise is one of the factors influencing the overall PAP size effects [12, 15]. Differently from the studies of Turner et al. [21] and Maloney et al. [17] in the present study, a weighted plyometric condition was not planned, which makes difficult to assess its influence on the size of PAP effects on deceleration performance and the recovery time needed.

Since the plyometric conditioning actions have kinematic similarities to explosive subsequent match activities, it has been suggested that they are more likely to specifically activate the higher order motor units (type II) associated with the following activity [12, 17]. In addition to the proposed mechanisms responsible for PAP previously introduced, Maloney et al. [17] suggested that an acute augmentation in limb musculotendinous stiffness, thanks to a plyometric activity, may contribute to eliciting PAP. In fact, the authors proposed that the active component of the muscle may benefit from the augmented stiffness increasing the force development, while the passive component may benefit from a higher elastic recoil. In this regard, Young and Elliott [26] stated that despite an augmented musculotendinous compliance would result in a higher storage and release of elastic energy, a stiffer system may assure a minimal delay between the stretching and the shortening phases of a stretch-shortening cycle (SSC), producing a good explosive performance, which is required in fast SSC (< 250 ms) such as sprinting. Therefore, in their review, Maloney et al. [17] stated that a stiffer system may enhance power activities, but only until the athlete’s optimal value is reached, beyond which performance would be impaired.

According to many authors [16, 17, 21, 27], one further reason supporting the use of plyometric exercises in order to elicit PAP is their extremely easier applicability in pre-competitive situations, compared to heavy resistance exercise, which require more time or equipment. The plyometric protocol used in this study does not require any equipment and can be easily performed in common spaces, both indoor and outdoor. To reinforce this idea, it should be highlighted that 75 seconds of exercise without any additional equipment led to a ∼ 4 % performance improvement. By looking at our data, it is interesting to note that in P condition, despite the immediate impairment of the performance (Δ time = + 0.31 %) and following the greatest improvement obtained at 2 minutes from the preload stimulus (Δ time = - 4.13 %), the enhancing-performance effects have gradually decreased over time (Figure 3, Table 1). This phenomenon is in accord to what expressed by the Maloney et al.’s review [17], that is the performance is impaired by the preload stimulus at the beginning, then improves thanks to PAP until a peak is reached and then decreases in an inverted U fashion. About this, Sale [14, 28] had already proposed that the longer the recovery time between the conditioning stimulus and the performance, the greater the recovery from fatigue, but also the greater the decay of PAP’s enhancing-effects.

Another interesting finding of this study is that at all time periods the trials performed in C condition resulted faster than Plyometric conditions (P) (Figure 5). This fact may be explained by a possible learning-effect that our participants could have experienced about the deceleration test, even if a familiarization session was previously scheduled in order to avoid such a phenomenon. Indeed, participants firstly executed the Plyometric condition, and on the following session day observed Control one. However, this data does not undermine the main findings of this study, given that performance improvements compared to the relative baseline have been observed only in the Plyometric condition (P), suggesting that the plyometric stimulus is responsible for the performance enhancements. Moreover, the performances of Plyometric and Control conditions were not similar since the baseline tests (Figure 5).

According to many authors [12, 17, 18, 21] a PAP may be evoked at the end of the warm-up to improve the following explosive activity after an individual adequate recovery. Furthermore, several authors suggested how a PAP can be elicited also in a training scenario to enhance the training stimulus of a following power exercise, in the so-called complex training strategy [11, 12, 15, 16, 17]. However, there are few evidences supporting the idea that PAP used in the complex training scenario could represent a more efficient training strategy than traditional ones [11, 16, 17].

Since PAP-related improvements have been reported for acceleration performance [20, 21] and along with this study for deceleration performance. For future researches it appears interesting to investigate possible effects of PAP on more complex tasks and more similar activities to those found in match situations, such as the change of direction ability, which is composed by accelerations and decelerations. Furthermore, sport performances like soccer are not determined by only physical factors (physiological and biomechanical), but it is strongly determined by cognitive, tactical and mental factors too [29]. Since all these factors are deeply connected, it appears interesting for future researches to investigate possible enhancing-performance PAP effects on more complex tasks like agility, defined as “*a rapid whole-body movement with change of velocity or direction in response to a stimulus*” by Sheppard and Young [8]. In fact, the athlete of situational team sports has to continuously adapt his activity in response to the surrounding situation.

Finally, although the potential beneficial effects of PAP to acutely improve short-term explosive performances such as jumps and sprints is clear, the influence that it may have on intermittent activities of team sports is not [15, 17]. According to Maloney et al. [17], the challenge would be evoking a potentiated state due to PAP and then maintaining it for the entire following duration of performance. Indeed, just letting the athlete start his performance in a potentiated state would provide him an advantage to start his performance [17]. In this sense, it has been proposed that in an intermittent and prolonged performance such as soccer, the series of contractions may act as conditioning stimuli themselves and so have a cumulative effect in eliciting PAP [14], along with an inevitable fatigue’s increase [17].

In conclusion, this study shows that a plyometric protocol of alternate-leg bound may enhance the deceleration performance at 2 minutes from its execution. Since acceleration and deceleration performance improvements have now been reported, it appears interesting for future research to investigate the possible effects of PAP on more complex tasks such as change of directions and agility, which are more similar to real sport activities.

## Acknowledgments

Open access funding provided by University of Vienna.

